# Validation of an automated segmentation algorithm for lower leg MR images, applied to sodium quantification

**DOI:** 10.1101/348136

**Authors:** Jasmine M. Greer, Ping Wang, Serpil Muge Deger, Aseel Alsouqi, T. Alp Ikizler, Jens M. Titze, Baxter P. Rogers

**Author notes:** Corresponding author (JMG). Current Address: Department of Radiology and Radiological Sciences, Vanderbilt University Institute of Imaging Science, Nashville, Tennessee, United States of America (Jasmine M. Greer). These authors contributed equally to this work (Jasmine M. Greer, Baxter P. Rogers). These authors contributed equally to this work (Ping Wang, Serpil Muge Deger, Aseel Alsouqi, T. Alp Ikizler, Jens M. Titze).

## Abstract

**Objective:** To develop and validate an automated segmentation algorithm for the lower leg using a multi-parametric magnetic resonance imaging protocol.

**Methods:** An automated algorithm combining active contour and intensity-based thresholding methods was developed to identify skin and muscle regions from proton Dixon MR images of the lower leg. Tissue sodium concentration was then computed using contemporaneously acquired sodium images with calibrated phantoms in the field of view. Resulting sodium concentration measurements were compared to a gold standard manual segmentation in 126 scans.

**Results:** Most cases had no observable errors in segmentation of muscle and skin. Six cases had minor errors that were not expected to affect quantification; in the worst, 126 mm^2^ (2%) of a muscle area of 8,042 mm^2^ was misclassified. In one case the algorithm failed to separate the tibia from the muscle compartment. Correlation between automated and manual measurements of sodium concentration was R^2^ = 0.84 for skin, R^2^ = 0.99 for muscle. Additionally, the RMSE was 2.4mM for skin and 0.5mM for muscle; the observed physiological range was 8.5 to 37.4mM.

**Conclusion:** For the purpose of estimating sodium concentrations in muscle and skin compartments, the automated segmentations provided equally accurate results compared to the more time-intensive manual segmentations. Sodium quantification serves as a biomarker for disease progression, which would assist with early diagnostic treatments. The proposed algorithm will improve workflow, reproducibility, and consistency in such studies.

## Introduction

With technological advancements, biomedical application of sodium (Na) magnetic resonance imaging is on the rise, as it provides unique and quantitative biochemical information related to tissue viability, cell integrity and function [1–7]. The lower leg muscle and skin is of particular interest because of the technical simplicity and speed of obtaining an MRI scan of the calf [8].

A straightforward approach to determine sodium levels based on the sodium magnetic resonance images of the calf is to manually segment the desired regions. This process requires meticulous attention and inconsistencies can be introduced via human error, which diminishes reproducibility.

An automated approach could address these problems, and consequently improve workflow. A variety of automated methodologies have been developed to segment anatomical magnetic resonance images of the leg. One approach has been to apply a fuzzy clustering method to segment anatomical regions such as adipose tissue, cortical bone, and spongy bone in the lower musculature of the leg [9] and in the thigh [10]. For segmentation, other studies use a combination of applications including: shaping histograms, adaptive thresholding, connectivity [11], a deformable model, global histogram based intensity thresholding, k means clustering [12], and intensity based temporal homomorphic filter [13].

In this study, we develop an application-specific automated segmentation pipeline for the lower leg and show that its segmentations applied to sodium MR images yield sodium concentration measurements comparable to the measurements obtained via the gold standard manual segmentations.

## Materials and Methods

### Participants

We conducted a study of 93 people who had formerly participated in a variety of sodium MRI studies in Vanderbilt University Medical Center between July 2014 and May 2017 and had data available with ^23^NaMRI readings. Our study sample included pre-hypertensive patients, maintenance hemodialysis patients, maintenance peritoneal dialysis patients, and controls. 31 people were scanned twice on separate occasions, while the remaining population was scanned once, in total 126 scans were acquired. The Institutional Review Board approved the study protocol and written informed consent was obtained from all study patients. The procedures were in accordance with the Declaration of Helsinki Principles regarding ethics of human research.

### MR Imaging

MR images were acquired on a Philips Achieva 3.0T MR scanner (Philips Healthcare, Cleveland OH, USA) using a ^23^Na quadrature knee coil (Rapid Biomedical GmbH, Rimpar, Germany). The left lower leg was placed in the coil, in close proximity to a set of calibration phantoms (NaCl aqueous solution of 10mM, 20mM, 30mM, and 40mM). Two proton scans were performed using the scanner body coil: a mDixon scan for fat and water images, and a standard proton-density-weighted image. These proton scans have the same geometry parameters: FOV = 192 x 192 mm^2^, resolution = 1 x 1 mm^2^, 5 slices at a thickness of 6 mm. The proton mDixon scan was acquired with TR = 200 ms and TE = 4.6 ms, 20 images were constructed in the form of water, water fat in-phase, water fat out-of-phase, and fat images, scan time = 3 min 52 s. The standard proton-density-weighted scan used the following parameters: TR/TE/FA = 4000 ms/ 30 ms/ 90°, and scan time = 2 min 32 s. Using the sodium coil and an optimized 3D gradient-echo sequence, a sodium image was obtained with the following parameters: FOV = 192 x 192 x 210 mm^3^, voxel size = 3 x 3 x 30 mm^3^, 7 slices, TR/TE/FA = 130 ms/0.99 ms/90°, bandwidth = 434 Hz/pixel, acquisition: 26, and scan time = 15 min 54 s [5].

### Manual Segmentation

Manual segmentation followed a previously described protocol [5]. The central imaging slices of the mDixon and sodium scans were used for manual segmentation. Five muscle regions of interest (anterior compartment, peroneus, soleus, medial gastrocnemius, and lateral gastrocnemius) were drawn on the mDixon, while a small region of the skin and phantoms were drawn on the sodium image (Fig. 1).

**Fig 1:**
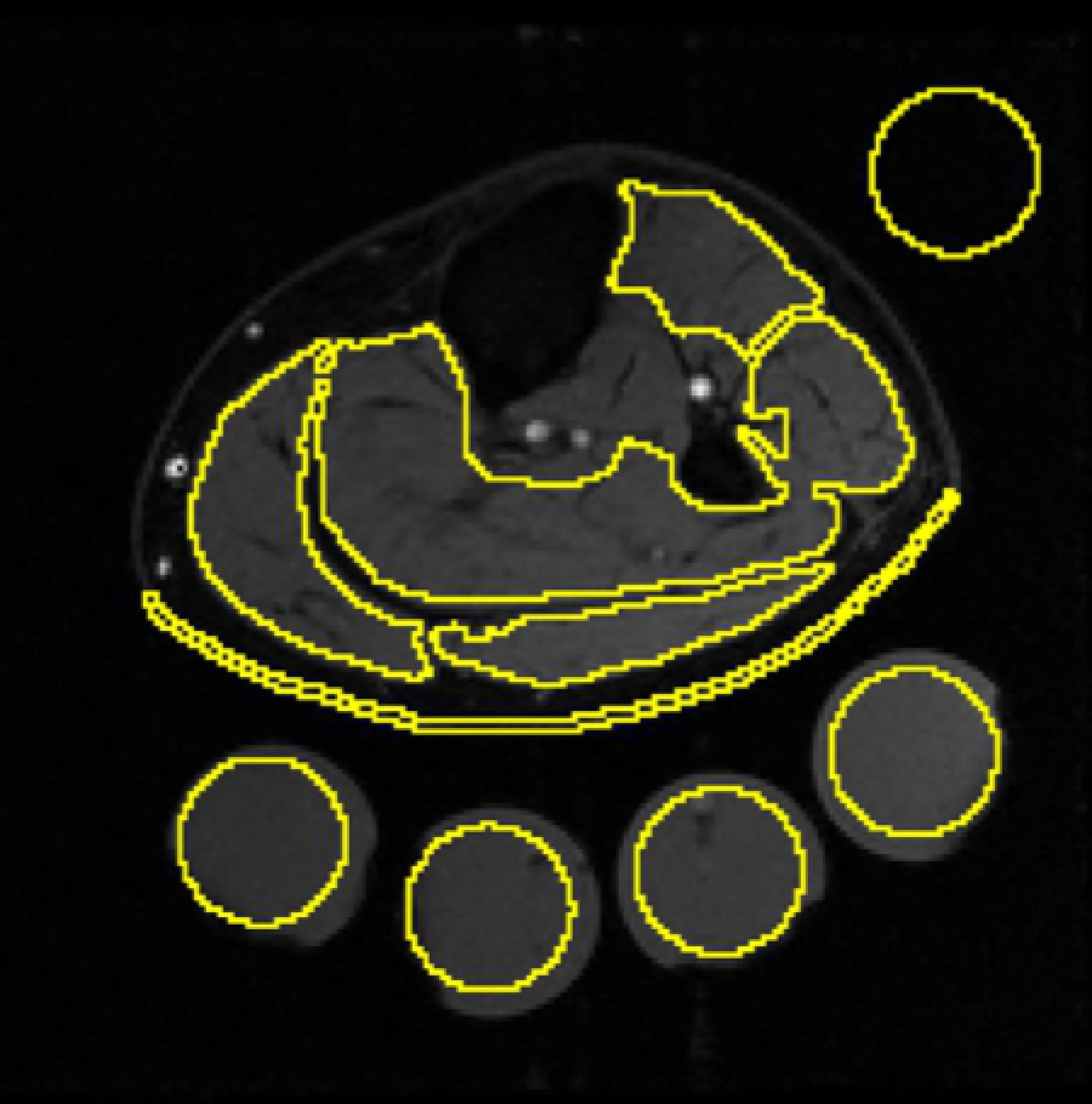
Example of manual segmentation. Four phantoms with sodium concentrations of 10mM, 20mM, 30mM, and 40mM, a background sample, skin, and 5 muscle regions were overlaid on a water image.

### Automated Segmentation

Image analysis was performed in MATLAB version 2016a (Mathworks, Natick, MA) using an XNAT data management platform [14].

Regions of interest were identified by applying an active contour model and a global histogram based intensity thresholding method. Active contour is an energy minimizing model that uses deformable curves to match the desired object [15]. First, the edges of the image are determined via application of

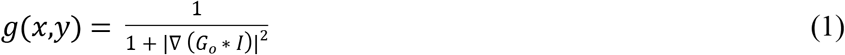
where “G_0_” represents the image and “I” represents the smoothing factor. The curves that shape the object are then minimized in order to closely identify the desired object. This is achieved by integrating the edge indicator function using calculus of variance

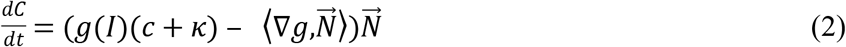

which yields a mask of the desired object. More simply stated, the active contour model can be thought of as creating a basic shape that encloses the object, then progressively moving closer to the object until it reaches the edge, shaping the object’s boundary (Fig 2).

**Fig 2.**
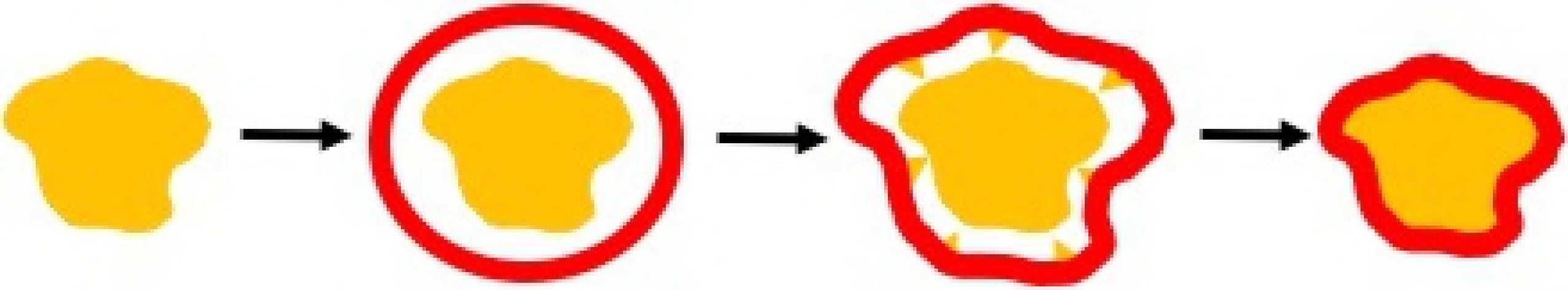
Basic illustration of the active contour method. A stepwise framework to illustrate the process of identifying an object outline.

The global histogram based intensity thresholding (Otsu’s method) is also used in this algorithm [16]. This method identifies a threshold that separates a bimodal intensity based histogram into two classes. In order to determine this threshold, the global histogram based intensity thresholding algorithm determines the thresholding that yields the smallest weighted variance of the two classes,

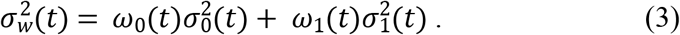

As illustrated in Fig 3, this thresholding approach distinguishes between the two groups of a bimodal histogram in order to produce a binary object.

**Fig 3.**
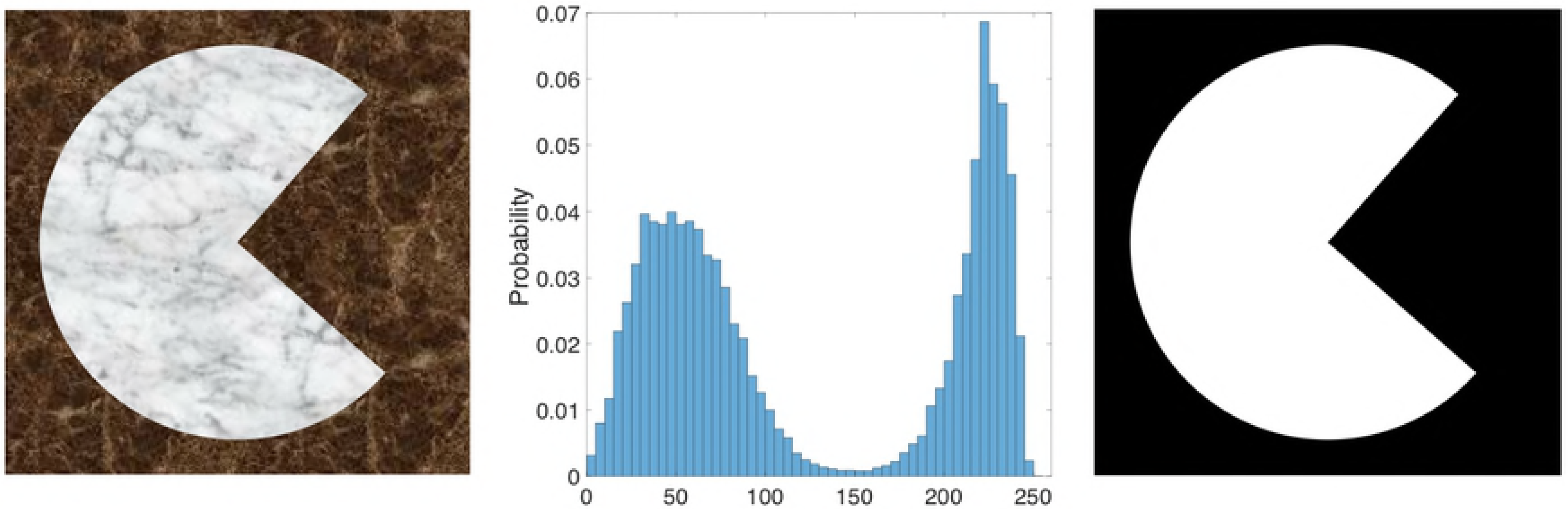
Basic illustration of global histogram based intensity thresholding. A water-only image derived from the mDixon scan was applied to an image intensity histogram to create two classes.

The automated quantification algorithm can be divided into four main phases: leg and phantom segmentation, skin segmentation, muscle segmentation, and quantification of sodium concentration.

First, using the proton-density weighted image (Fig 4a) a 400-iteration active contour Chan-Vese method [15] was used to identify the leg portion of the mask and phantoms from the background (Fig. 4b). Nature of each segmented region was determined automatically based on size (leg >2400 mm^2^, phantoms <1300 mm^2^).

**Fig 4.**
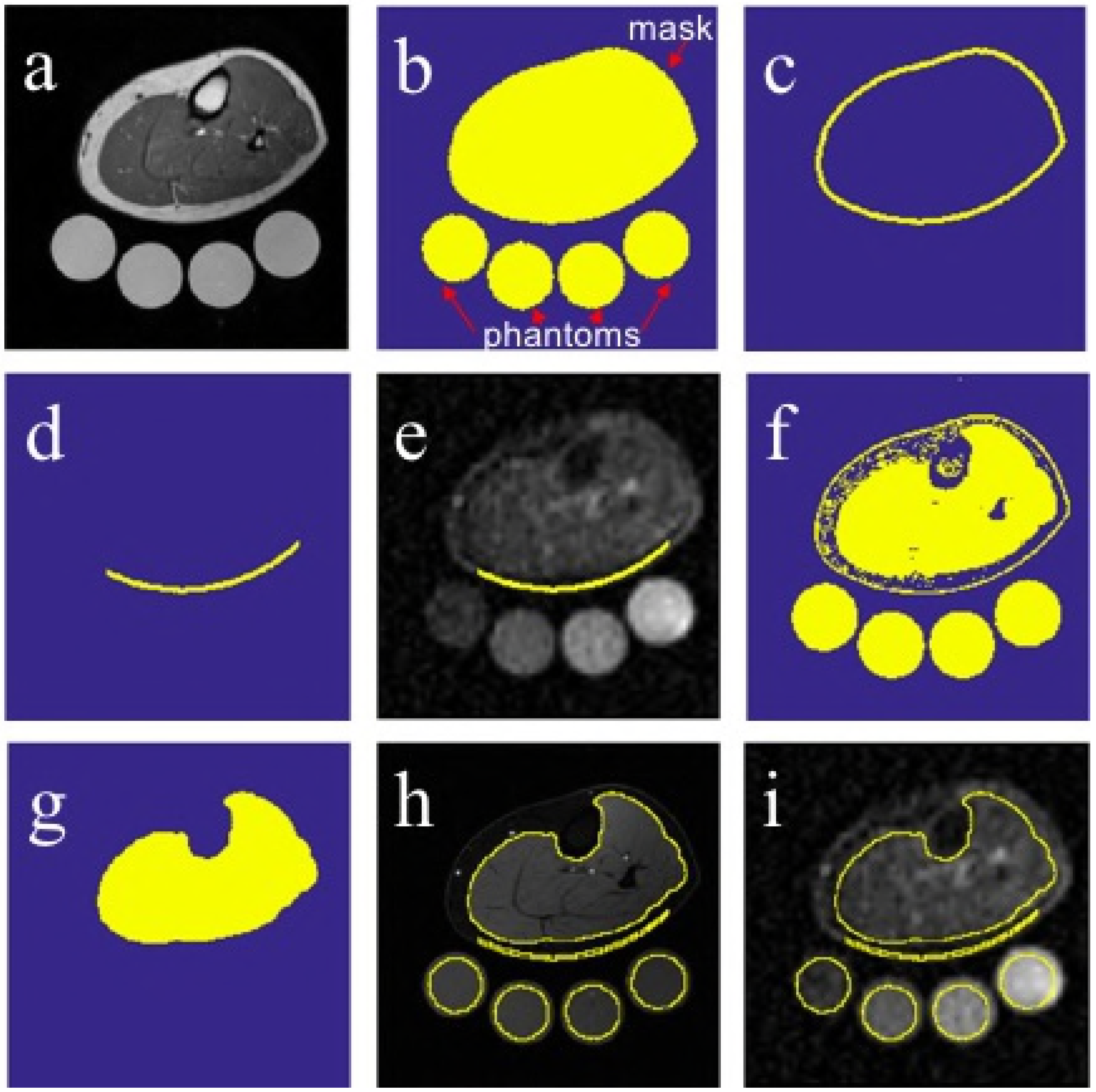
Creation and application of mask from MR images. (A) Proton density image, (B) mask of leg and phantom regions, (C) mask of skin region, (D) reduced skin mask, (E) reduced skin mask overlaid on sodium image, (F) Otsu thresholding of the water image, (G) smoothed muscle region, (H) phantom, skin, and muscle regions overlaid on water image, and (I) phantom, skin, and muscle regions overlaid on sodium image.

The skin region was estimated by eroding the leg portion of the mask (Fig. 4b) by a 4 mm radius circular kernel and subtracting the resulting image from the original leg portion of the mask (Fig. 4b) to select approximately the outer 4 mm of the leg region (Fig. 4c). It should be noted that this process assumes the skin thickness is similar in all participants. At the time of MR acquisition, the posterior area of the leg was resting on the phantom holder surface and thus aligned perpendicular to the slice direction such that through-plane partial volume effect was minimized [22]. Therefore, the skin region was reduced to include only the portion in contact with the surface of the phantom holder (Fig. 4d). The reduced skin region was parallel to the coil surface and the tissue thickness was more stable. Then, the produced image was overlaid on the sodium image (Fig. 4e) [17, 18].

The muscle region was identified on the water-only image derived from the mDixon scan (Fig. 3a) using a two-class global histogram based intensity thresholding method (Fig. 3) [19]. The estimated classification threshold was then reduced by 50% to account for intensity inhomogeneity in the image. Both the resulting two class image (Fig. 4f) and the leg portion of the mask (Fig. 4b) which was eroded by a 2 mm radius circular kernel, were used to create a three class intensity based leg image. By utilizing the index values for identification purposes, the region that included the muscle was isolated. Following erosion by a 1 mm radius circular kernel, size based artifact removal, and dilation by a 1 mm radius circular kernel, all remain holes within the muscle region were filled. The extracted muscle region was smoothed using a 300 iteration Chan-Vese active contour model with a smooth parameter of 1.2. The skin region (Fig. 4c) was then subtracted from this muscle region to confirm there is no overlap between the two (Fig. 4g). Then, the four phantoms were uniquely labeled and eroded by a 4 mm radius circular kernel to ensure proper alignment on the sodium image. The phantoms, skin, and muscle regions are shown overlaid on the water only image in Figure 4h [17, 18].

### Calibration for Quantitative Sodium Concentration

To quantify the sodium content in each region, the linear relationship between tissue intensity and the calibration phantoms was applied. A linear fit was computed using the known concentrations of the phantoms: 10mM, 20mM, 30mM, and 40mM, and their respective average sodium image intensity signal to estimate the calibration coefficients. Using these parameters, the sodium image was calibrated. The regions of interest were then applied to this calibrated sodium image (Fig. 4i) and the mean and median sodium concentrations were quantified [17, 18].

## Results

### Participants and Scan Quality

In total, 126 scans were acquired from 93 participants. Three scans were excluded from the analysis based on technical errors at the time of acquisition: in one case, the phantom holder was misaligned relative to the leg, while in two cases, the fields of view were misaligned between sodium and proton scans. Method comparisons were based on the remaining 123 scans.

### Automatic Segmentation Algorithm Results

Of the 123 usable scans, 94% of the segmentations were highly accurate on visual inspection, correctly identifying the muscle and skin while excluding the tibia (Fig. 4h-i). 5% were usable for the intended purpose, identifying the muscle and skin with minor errors and excluding the tibia (Fig. 5a). We observed that two of the cases with minor errors also had high amounts of intramuscular fat, e.g. Fig. 5b. And lastly, there was one case in which the tibia was not excluded (Fig. 5c).

**Fig 5:**
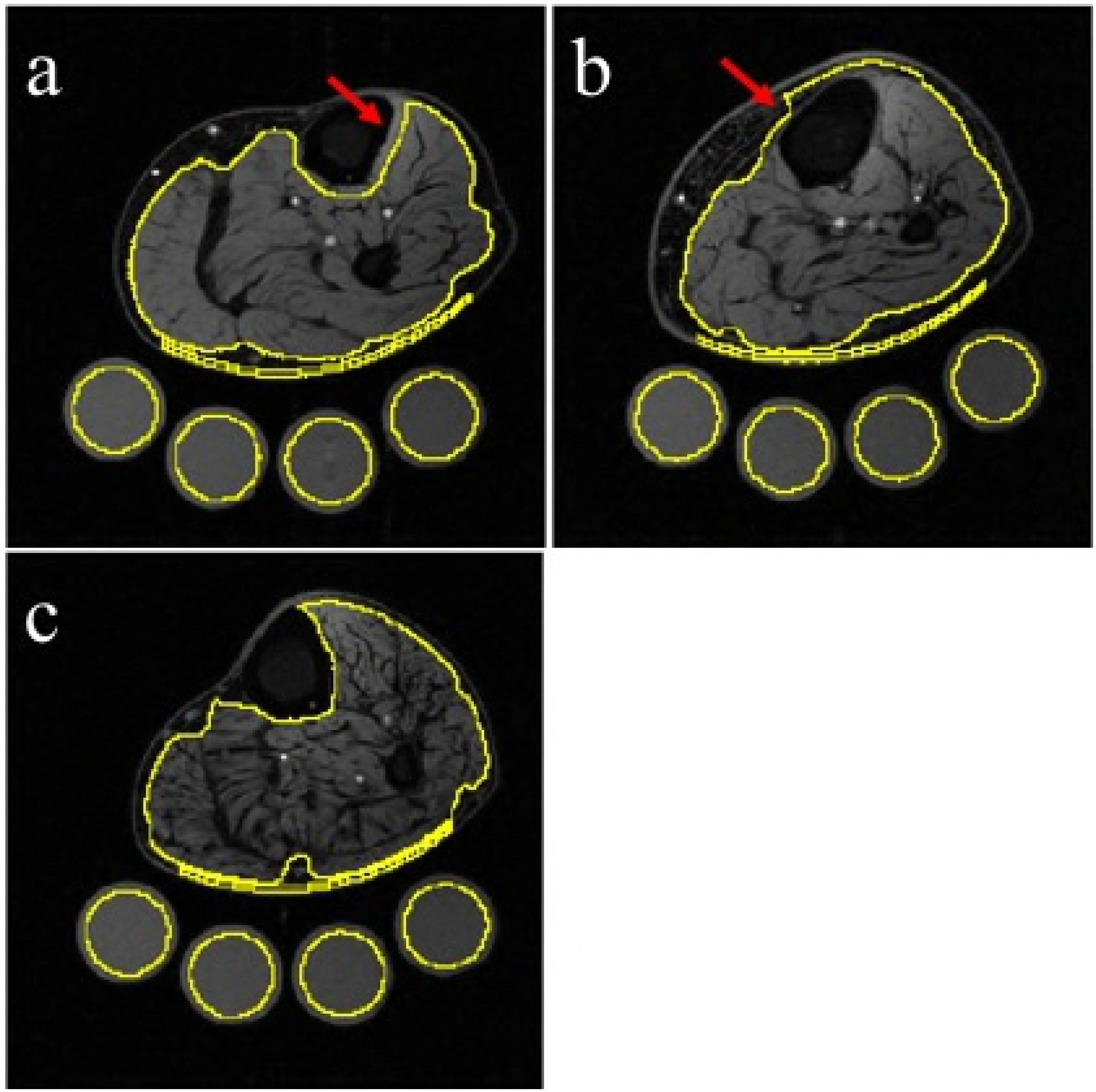
Illustration of automated segmentations special cases. (A) Example of a minor error, exclusion of small amount of muscle tissue (arrow). (B) In one case, the tibia was erroneously included in the muscle region. (C) A case with high level of intramuscular fat.

### Automatic and Manual Segmentation Comparison

Sodium concentrations estimated by the automated algorithm were compared to values obtained from the gold standard manual segmentation using Bland-Altman analysis (Fig. 6) [20]. The bias for both regions was approximately zero: -0.05 mM for the skin and -0.11 mM for the muscle. The root mean square error (RMSE) of the automated algorithm compared to gold standard was 0.5 mM for the muscle, and 2.4 mM for the skin. The range of sodium concentration values in the muscle and skin regions in the entire sample was 11.3 to 35.0 mM (muscle) and 8.5 to 37.4 mM (skin). The correlations between automated and manual measurements were 0.99 and 0.84 for the muscle and skin, respectively. We observed four cases in the skin region marked in red on Fig. 6a) and three cases in the muscle region (marked in red on Fig 6b) where the difference between automated and manual measurements was more than two standard deviations from the mean.

**Fig 6.**
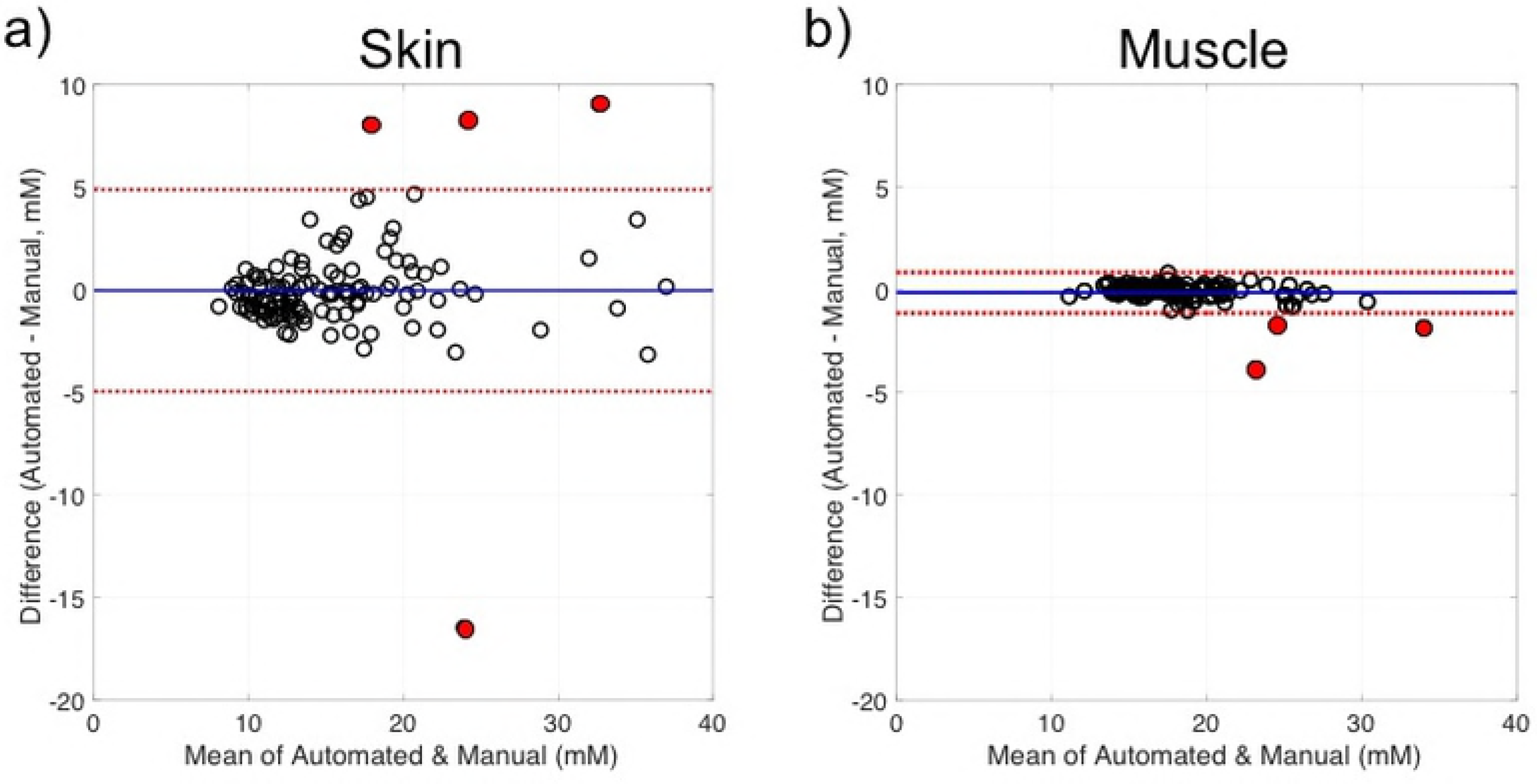
Bland-Altman plots comparing automated and manual measurements of sodium concentration in the skin and muscle. (A) Bland-Altman plot of the skin comparing the automated and manual measurement differences (y axis) versus the automated and manual measurement mean (x axis) (sodium range: 8.5-37.4 mM). Discrepancy cases indicated in red. (B) Bland-Altman plot of the muscle, comparing the automated and manual measurement differences (y axis) versus the automated and manual measurement mean (x axis) (sodium range: 11.3-35.0 mM). Discrepancy cases indicated in red.

### Automatic and Manual Segmentation Inter-Scan Comparison

Of the 123 used scans, 31 subjects were scanned twice. Using the Bland-Altman analysis (Fig6) [20], the inter-scan comparison was evaluated. The bias for both methods and both regions were approximately zero. In all cases, bias was below 1.05 and 95% limits of agreement were less than +/−10 mM. The limits of agreement for the automated method of the muscle was [-4.50, 4.87], the manual method of the muscle was [-4.91, 5.44], the automated method of the skin was [-4.87, 5.12], and the manual method of the skin was [-5.91, 8.01].

## Discussion

In this study, we aimed to develop an algorithm that would allow us to streamline ^23^NaMRI readings of the lower leg. Our data suggest that the sodium concentration measurements obtained by the automated segmentation were of excellent quality, adequate to replace those obtained by the gold standard manual segmentation method.

Seven cases fell outside the limits of agreement in the Bland-Altman analysis, indicating that these cases had a relatively large discrepancy between manual and automated results: four were in skin and three cases in muscle. For the manual segmentation of the skin region, we observed variability from one case to the next in how much skin versus background was included in the final region, and in the thickness of the manually drawn skin region. This is a challenging segmentation task even when the regions are directly drawn on the sodium image [5], because the skin (~2 mm thick) is poorly resolved at the 3 x 3 mm in-plane voxel size. The automated method sacrificed any improvement in accuracy related to using the sodium image intensity to define boundaries; however, it added substantial consistency in positioning and thickness due to the use of the higher resolution structural images. Three of the four cases where skin results were outside the limits of agreement (Fig. 6a) showed erroneous inclusion of background voxels or exclusion of skin voxels in the manual segmentation. In the fourth case, the source of inconsistency was unclear.

Based on the Bland-Altman analysis of the muscle (Fig. 6b) the sodium concentration measurements in this region were highly correlated. However, three cases fell outside of the limit of agreement. One of these had high levels of intramuscular fat (Fig. 5b). In this scenario, the manual approach which divides the muscles into five sub-compartments could possibly exclude slightly more of the intramuscular fat tissue between compartments compared to the automated method, resulting in a small bias towards lower muscle sodium concentration estimates by the automated method. Another was the case where the tibia was erroneously included in the muscle region due to poor estimation from the active contour model in the automated method, this consequently yielded an underestimation of sodium concentrations. The manual segmentation from the third case was performed on the first slice of the mDixon scan instead of the middle slice, which was an accidental deviation from the protocol due to human error as is typical with manual procedures.

Overall, the inter-scan comparisons were comparable for both the regions and segmentation methods (Fig. 7). The correlations between baseline and the follow up for the skin region is slightly higher for the automated method, hence increased reproducibility, compared to the manual method. Conversely, the correlations between baseline and the follow up for the muscle region is the same in the automated and manual methods. It should be noted that the used subdataset is from an ongoing longitudinal study during which some subjects will experience treatment and time effects, thus we are not strictly measuring reproducibility.

**Fig 7.**
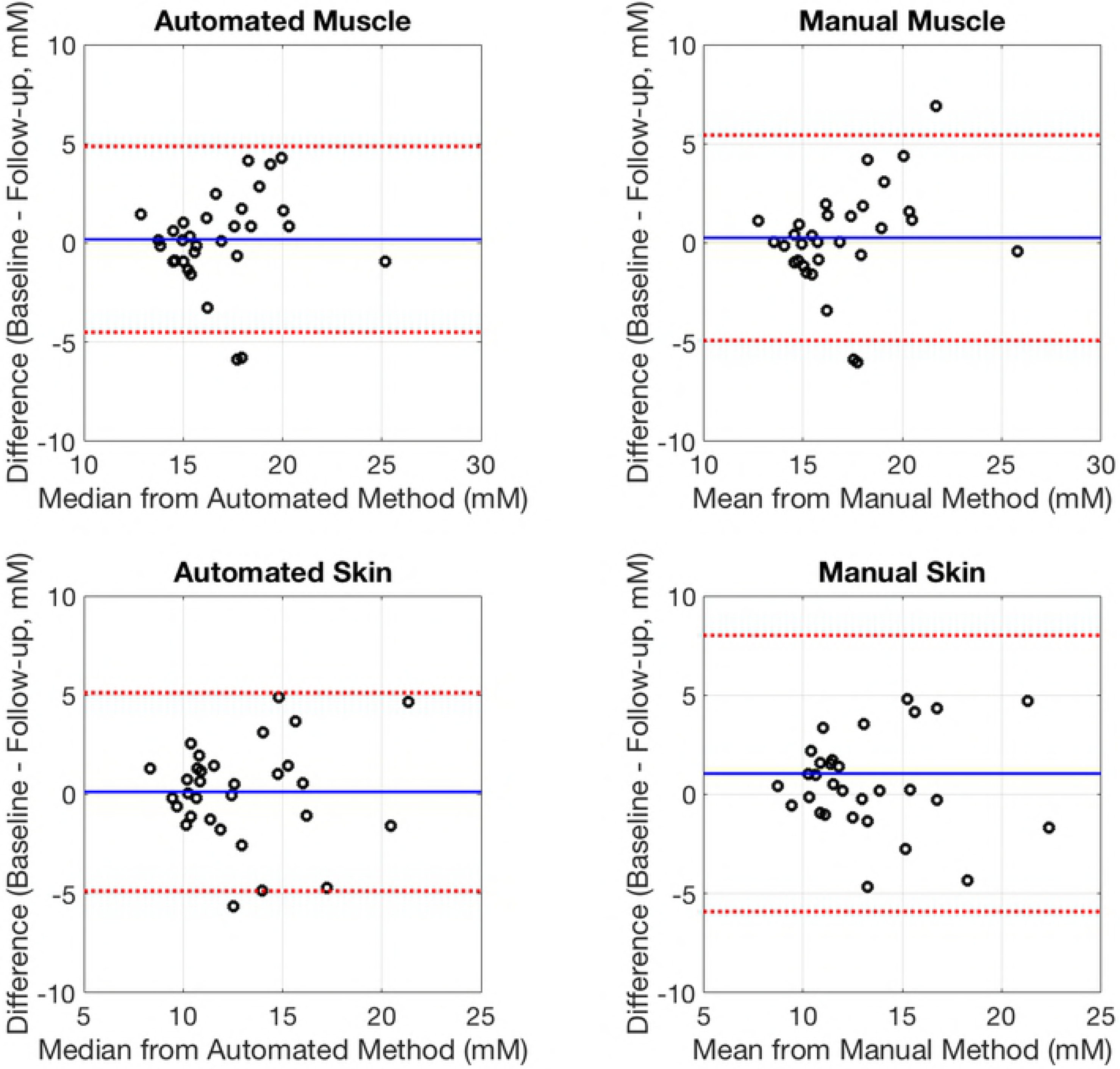
Bland-Altman plots comparing sodium concentration measurement concentration in the skin and muscle before and after the study. (A) Bland-Altman plot of the muscle region, identified by the automated method, comparing the baseline and follow-up measurement differences (y axis) versus the sodium median from the automated method (x axis) (sodium range: 12.9-25.2 mM). (B) Bland-Altman plot of the muscle region, identified by the manual method, comparing the baseline and follow-up measurement differences (y axis) versus the sodium mean from the manual method (x axis) (sodium range: 12.7-25.8 mM). (C) Bland-Altman plot of the skin region, identified by the automated method, comparing the baseline and follow-up measurement differences (y axis) versus the sodium median from the automated method (x axis) (sodium range: 8.32-21.3 mM). (D) Bland-Altman plot of the skin region, identified by the manual method, comparing the baseline and follow-up measurement differences (y axis) versus the sodium mean from the manual method (x axis) (sodium range: 8.72-24.8 mM).

Although the automated algorithm performed quite well, there are some limitations. First, the MR imaging protocol it relies on is fast, but still moderately complex and multi-parametric, requiring both proton-tuned and sodium-tuned coils. Also, the results still require manual quality review to identify significant failures such as inclusion of the tibia. Finally, the algorithm did not reliably exclude the fibula from the muscle region. However, the area of the fibula is relatively quite small, and we used the median instead of the mean in the muscle region to summarize sodium quantification more robustly in the presence of a small number of outlier voxels (fibula). Observed results had trivial errors compared to the manual segmentation that consistently excluded the fibula (Fig. 7).

The global intensity-based thresholding step can be confounded by intensity inhomogeneity in the MR images, resulting in a threshold that excludes some muscle tissue in lower intensity regions or includes some background voxels in higher intensity regions. Such inhomogeneity was present in our data, but not to a degree that affected results compared to manual segmentation. This issue could be more pronounced in data from other field strengths or other imaging protocols. Possible solutions would be to utilize a bias field correction [21] or a local thresholding method so that the intensity-based segmentation is applied more uniformly across the image.

Finally, arbitrary fixed smoothness parameters were chosen for the active contour portion of the segmentation algorithm, which is a likely cause of the minor errors in identifying the muscle edges. Developing a dynamic and automated procedure for tuning smoothness to match specific images may improve results, although we would expect the improvement to be minimal in terms of the final concentration measurements.

The algorithm is applied to the proton Dixon image. The resulting segmentation in this case were applied to a standard sodium weighted image, but in general could be applied to images of other physiological parameters. For instance, intracellular or extracellular sodium concentrations can be measured separately using inversion recovery techniques [1]. Additionally, to obtain more accurate sodium concentration measurements in the skin region, ^23^NaMRI at 7.0 T could be used to acquire higher image resolution [22].

## Conclusion

In summary, we developed an algorithm that could streamline assessment of ^23^NaMRI measurements in the leg in both research and clinical settings. By applying an active contour model and a global histogram based intensity thresholding method, specific regions of interest were identified which were later used for sodium quantification. This algorithm proved to be highly comparable to the gold manual segmentation method. For both the skin and muscle regions, the RMSE was relatively low based on the physiological range and the bias was approximately zero based on the Bland-Altman analysis. This automated approach is time efficient, reproducible, and minimizes observer bias and human error. Our results suggest that this algorithm is an excellent alternative to the manual segmentation methodology.

## Acknowledgements

This work was conducted in part using the resources of the Advanced Computing Center for Research and Education at Vanderbilt University, Nashville, TN and the resources of the Center for Computational Imaging at Vanderbilt University Institute of Imaging Science, Nashville, TN. We would also like to acknowledge the Human Imaging Core at Vanderbilt University Institute of Imaging Science for their support.

